# Geometrically-engineered organoid units and their assembly for pre-construction of organ structures

**DOI:** 10.1101/2024.05.06.592718

**Authors:** Ayaka Kadotani, Gen Hayase, Daisuke Yoshino

**Author notes:** **Corresponding author (Lead author):** (Daisuke Yoshino, Ph.D.). **Co-corresponding author:** (Gen Hayase, Ph.D.).

## Abstract

Regenerative medicine is moving from the nascent to the transitional stage as researchers are actively engaged in creating mini-organs from pluripotent stem cells to construct artificial models of physiological and pathological conditions. Currently, mini-organs can express higher-order functions, but their size is limited to the order of a few millimeters. Therefore, one of the ultimate goals of regenerative medicine, “organ replication and transplantation with organoid,” remains a major obstacle. 3D bioprinting technology is expected to be an innovative breakthrough in this field, but various issues have been raised, such as cell damage, versatility of bioink, and printing time. In this study, we established a method for fabricating, connecting, and assembling organoid units of various shapes independent of cell type, extracellular matrix, and adhesive composition (unit construction method). We also fabricated kidney tissue-like structures using three types of parenchymal and interstitial cells that compose the human kidney and obtained findings suggesting the possibility of crosstalk between the units. We anticipate that this technique will bring us closer to the realization of highly efficient and rapid fabrication of full-scale organoids that can withstand organ transplantation.

## INTRODUCTION

The development of three-dimensional (3D) culture systems, organoids, has been the most exciting advance in medical and biological fields in the last two decades since the invention of iPS cells [1–6]. This organoid system has enabled the modeling of genetic, degenerative, and cancer diseases that were difficult to reproduce *in vitro* [7–11], and has been expected to lead to innovative advances in establishing medical treatments. Organoids can express the higher-order functions of organs [12], but are currently limited to a few millimeters in size, referred to as “mini-organs” are called. Therefore, one of the ultimate goals of regenerative medicine, “organ replication and transplantation using organoids,” faces a major obstacle.

Currently, 3D bioprinting is the most promising method for producing artificial tissue that can be used as organs for transplantation. Extrusion-based bioprinting (EBB), in which a solution containing living cells, bioink, is extruded and layered in a 3D space such as a 3D printer, is a mainstream method [13]. EBB has the advantage of being able to form complex structures, including internal architecture, without the need to prepare molds with cell-dense solutions as bioinks [14]. Conversely, problems with EBB include cellular damage due to pressure and shear forces during bioink ejection, the difficulty of developing bioinks with appropriate mechanical, structural, and biological properties, and the long time required to print full-size organs [15–17]. Volumetric Bioprinting (VBP), unlike conventional methods (*i.e.*, layer-by-layer stacking), creates an object by photo-crosslinking the resin by exposing it to a calculated two-dimensional (2D) light pattern while rotating a transparent container [18]. As a result, it takes only a few seconds to a few minutes to fabricate a model, regardless of size. The features of short fabrication time and nozzle-less fabrication do not affect cell viability and functionality and provide high resolution (up to tens of micrometers) [19]. Hence, VBP is a new and promising technique that overcomes several of EBB’s limitations. However, it is difficult to precisely control the position and density distribution of cells in the pre-cured gel with current methods [19, 20], and it is expected to be challenging to construct full-size organs composed of different cells and extracellular substrates. Therefore, it is desirable to develop a technique for artificially constructing organ-size 3D tissues with high efficiency and yield.

To address these current problems, a method of 3D tissue engineering using organoid building blocks (OBBs), *i.e.*, stacking OBBs such as spheroids and organoids to reproduce organ-specific functions, has been developed [21]. The stacking of organoids and spheroids ranging in size from several hundred µm^3^ to 1 mm^3^ is expected to significantly reduce the printing time currently required for the spatial arrangement of single cells, bringing us closer to the rapid construction of human tissues. In particular, the OBB has recently been introduced into a 3D bioprinter, enabling large-scale organoid-like structures, assembloid, by realizing rapid spatial arrangement [21–24]. Although this technology solves the heterogeneity of iPS cell-derived organoids, more is needed to address the limitations of the types of bioinks, the size of the OBBs that can be used, the complicated spatial arrangement due to the basic shape of the block, such as a sphere, and the changes in bioink properties associated with printing due to the use of a 3D bioprinter.

In contrast, we conceptualize establishing a method to assemble organoid units of various shapes pre-divided into basic organ elements like LEGO® (unit construction method). This leads to a highly efficient and rapid technique to produce full-scale organoids that can withstand organ transplantation. We envision the production of organoids with transplantable sizes in the future. In particular, we focus on the construction of full-scale kidney organoids as one of the treatment options for renal disorders, which are difficult to cure with drugs. Here, we show our method for fabricating organoid units with six different shapes and the cellular dynamics inside the units when stacked, as well as the results of forming and assembling the units using three types of parenchymal and interstitial cells that compose the human kidney.

## MATERIALS AND METHODS

### Cell culture

An easy-to-handle cancer cell line was used to validate the feasibility of fabricating the organoid units and their assembly. Green fluorescent protein (GFP)-labeled human breast adenocarcinoma cell line (MDA-MB-231; AKR-201, Cell Biolabs, San Diego, CA, USA) was cultured with Dulbecco’s modified eagle medium (DMEM; 31600-034, Gibco, Thermo Fisher Scientific, Waltham, MA, USA) containing 10% heat-inactivated fetal bovine serum (FBS; S1810, BioWest, Nuaillé, France) and 1% penicillin-streptomycin (P/S; 15140-122, Gibco). Three types of human primary cells were also used to validate the construction of organ-like structures by co-culturing parenchymal and interstitial cells: human renal glomerular epithelial cells (HRGEpCs; 942-05n, Cell Applications, San Diego, CA, USA), human glomerular mesangial cells (NHMCs; ACBRI127, Cell Systems, Kirkland, WA, USA), human umbilical vein endothelial cells (HUVECs; 200-05n, Cell Applications). The primary cells were cultured with Medium 199 (31100-035, Gibco) containing 5% FBS, 1% P/S, 10 µg/L human basic fibroblast growth factor (bFGF; GF-030-3, AUSTRAL Biologicals, San Ramon, CA, USA), 10 µg/L human epidermal growth factor (hEGF; E9644, Sigma-Aldrich, St. Louis, MO, USA), 1% ITS-X supplement (094-06761, Fujifilm Wako Pure Chemical Corp., Osaka, Japan), 36 µg/L hydrocortisone (50-23-7, MP Biomedicals, Irvine, CA, USA), and 4 ng/L 3,3′,5-triiodo-L-thyronine sodium salt (T6397, Sigma-Aldrich). The cells were cultured in a 75 cm^2^ flask (658175, Greiner Bio-One, Kremsmünster, Austria) pre-coated with/without 0.1% bovine gelatin solution (G9391, Sigma-Aldrich) until reaching 90% confluence. Primary cells from the fifth to ninth passages were used for experiments in this study.

### Mold processing

Monolithic porous bulk material with superhydrophobicity (boehmite nanofiber-polymethylsilsesquioxane; BNF-PMSQ) [25] was processed using a CNC milling machine (monoFab SRM-20 or MDX-50, Roland DG, Shizuoka, Japan) to manufacture molds for various shapes of organoids. We fabricated bead ring, cylinder, rectangular solid, cubic, and sheet-shaped molds in addition to the spherical one [25, 26] (**Supplementary Fig. 1**). The mold models were created in a 3D-CAD (SolidWorks, Dassault Systémes SOLIDWORKS Corp., Waltham, MA, USA) and then exported in STL format, from which the processing paths were coded by CAM software (SRP Player, Roland DG). Processing was performed in two stages, roughing and finishing, to prepare the surface of the monolithic porous material and make it superhydrophobic (**Supplementary Table 1**). Polytetrafluoroethylene (PTFE) was used as the mold material when necessary.

### Organoid unit fabrication

To validate the feasibility of fabricating the organoid units and their assembly, the MDA-MB-231 cells were harvested after reaching 90% confluence with 0.25% trypsin-EDTA (25200-072, Gibco) and resuspended in the culture medium at a concentration of 5.0 × 10^7^ cells/mL. Cell-suspended collagen solution [4.0 mg/mL; native collagen acidic solution (IAC-50, KOKEN, Tokyo, Japan), 10× DMEM, 10 mM NaHCO_3_, 10 mM HEPES-NaOH (pH7.5), and the cell suspension] was prepared on ice to give the final concentration of 5.0 × 10^6^ cells/mL. The cell-suspended collagen solution was then dispensed onto the sterilized molds in a predetermined amount and order (**Table 1**). The dispensed solution was allowed to stand still in a CO_2_ incubator (37°C in a 100% humidified atmosphere of 5% CO_2_) for 30–60 min. After gelation, the primary organoid block was transferred to a 35 mm diameter dish (3000-035, AGC Techno Glass, Shizuoka, Japan) by dropping a small amount of medium to cover it, poking it with the tip end of a micropipette to float it, and then adding more medium to pour it in. The units were incubated and matured for up to 5 days while being observed under a stereomicroscope (SZX16, Olympus, Tokyo, Japan) or a fluorescence imaging system (THUNDER Imaging System, Leica Microsystemes, Wetslar, Germany).

**Table 1.**
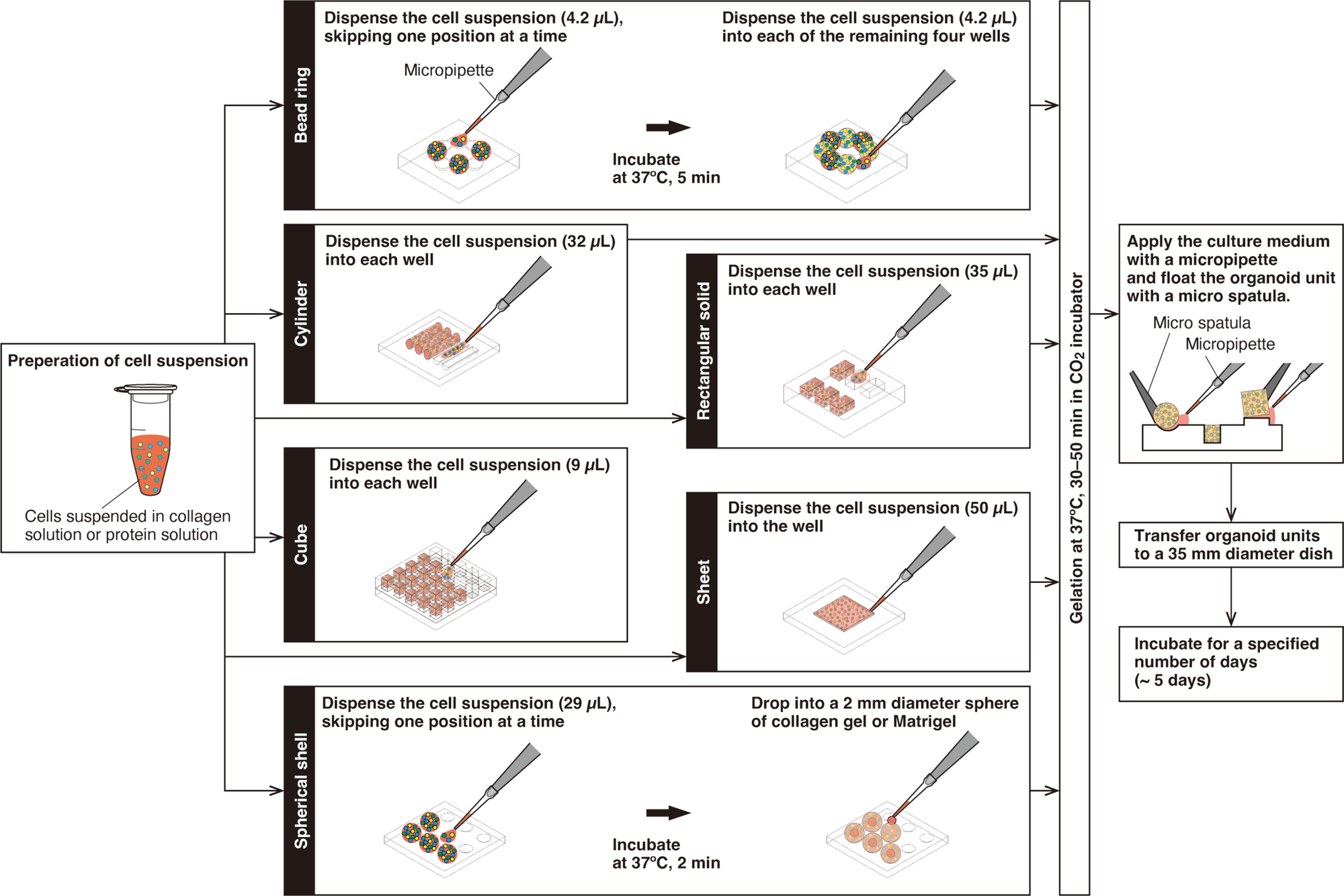
Procedure for fabrication of organoid units with each geometry.

To validate the construction of organ-like structures with the units in tri-culture, the HRGEpCs, NHMCs, and HUVECs were harvested after reaching 90% confluence with 0.05% trypsin-EDTA (25300-062, Gibco) and resuspended in the culture medium at a concentration of 5 × 10^8^ cells/mL. Cell-suspended protein solution [4.5 mg/mL Matrigel (Matrix for Organoid Culture, 356255, Corning, Corning, NY, USA), 1.5 mg/mL native collagen acidic solution, 10× DMEM, 10 mM NaHCO_3_, 10 mM HEPES-NaOH (pH7.5), and the cell suspensions] was prepared on ice to give the final concentrations of 0.5 × 10^7^ cells/mL (HRGEpCs and NHMCs) and 4.0 × 10^7^ cells/mL (HUVECs), respectively (*i.e.*, HRGEpCs:NHMCs:HUVECs=1:1:8). Tri-cultured organoid units, similar to MDA-MB-231 units, were formed and gelatinized with the prepared solution. They were then transferred to 48-well plates (VTC-48, AS ONE Corp., Osaka, Japan) and cultured for 7 days using OncoPro medium (A5701201, Gibco) supplemented with 1% P/S, 10 µg/L bFGF, and 10 µg/L hEGF.

### Organoid unit assembly (unit construction)

The units were first transferred to an empty tissue culture dish using a micro spatula (6-524-01, AS ONE Corp.) for block-to-block assembly. We applied 5-10 µL of adhesive to the area to be bonded with a micropipette, stuck the units together, and incubated for 2-5 minutes. Collagen solution [2.5 mg/mL; native collagen acidic solution, 10× DMEM, 10 mM NaHCO_3_, 10 mM HEPES-NaOH (pH7.5), and ultrapure water] was used as the adhesive for bonding MDA-MB-231 units, and protein solution [4.5 mg/mL Matrigel, 1.5 mg/mL native collagen acidic solution, 10× DMEM, 10 mM NaHCO_3_, 10 mM HEPES-NaOH (pH7.5), and the cell suspension (HUVECs; final concentration, 1.0 × 10^6^ cells/mL)] was used for bonding tri-cultured units. After bonding, the assembled units were transferred to the 35 mm diameter dish filled with the culture medium using a medicine spoon to continue incubation.

### Antibodies

The sheep polyclonal anti-Nephrin antibody (Cat# AF4269) was purchased from R&D Systems (Minneapolis, MN, USA). The mouse monoclonal anti-CD31 antibody (Cat# 3528) was purchased from Cell Signaling Technology (Danvers, MA, USA). The FITC-conjugated human monoclonal anti-CD90.1 antibody (Cat# 130-112-683) was purchased from Miltenyi Biotec (Bergisch Gladbach, Germany). Alexa Fluor 594-conjugated goat anti-mouse IgG (Cat# A-11032) and Alexa Fluor 647-conjugated donkey anti-sheep IgG (Cat# A-21448) secondary antibodies were purchased from Thermo Fisher Scientific.

### Immunohistochemistry

Cultured organoid units were fixed with 4% paraformaldehyde phosphate buffer saline (PFA; 163-20145, Fujifilm Wako Pure Chemical Corp.) for 1 h at room temperature (RT). The units were cryoprotected by soaking in 20% sucrose/phosphate buffered saline (PBS; 05913, Nissui Pharmaceutical, Tokyo, Japan) for 5 h and 30% sucrose/PBS for an additional overnight at 4°C. Fixed units were frozen in optical cutting temperature (OCT) compound (45833, Sakura Finetek Japan, Tokyo, Japan) and cut into 15 µm-thick frozen sections on cryofilm using a cryostat (CM1860, Leica Microsystems). After cutting out the frozen sections, the cells were permeabilized with 0.1% Triton X-100 (17-1315-01, Pharmacia Biotech, Uppsala, Sweden) in PBS, followed by incubation in 1% Block Ace (BA; UKB40, KAC, Kyoto, Japan) in PBS to prevent nonspecific antibody absorption. The cells were then stained using the primary and secondary antibodies diluted in 1% BA in PBS and PBS, respectively. Cell nuclei were stained using 4’,6-diamidino-2-phenylindole (DAPI; D1306, Invitrogen, Thermo Fisher Scientific). Stained organoid unit sections were observed with optical sectioning fluorescence microscopy (Axio Observer 7 with Apotome 3, Carl Zeiss, Oberkochen, Germany).

### Immunofluorescence staining

Cultured organoid units were fixed with 4% PFA for 1 h. The cells were permeabilized and blocked with 0.15% Triton-X100 and 1% BA in PBS for 1 h. The cell nuclei and actin cytoskeletal filaments were stained using DAPI and Alexa Fluor 594 Phalloidin (A12381, Invitrogen), respectively, for 1 h. At each step, the samples were washed with PBS for 5 min. After staining, the units were left to soak overnight in PBS at 4°C. For multiplex staining with antibodies, the samples were made transparent. After fixation, the units were immersed overnight in 50% Tissue-Clearing Regent CUBIC-L (T3740, Tokyo Chemical Industry, Tokyo, Japan) containing 500 mM NaCl, followed by membrane permeabilization and blocking for 1 h. The cells were stained overnight at 4°C with primary antibodies (diluted in 1% BA in PBS) and secondary antibodies (diluted in PBS), respectively, and then post-fixed with 4% PFA for 1 h. Cell nuclei were also stained with DAPI. The stained units were soaked in 50% Tissue-Cleaning Reagent CUBIC-R+(N) (T3983, Tokyo Chemical Industry) for 20 min and observed in 100% CUBIC-R+(N). Unless otherwise noted, the staining process was performed at RT, with three 15-minute washes with PBS between each step. All processes were carried out with shaking. Fluorescent images of the stained units were obtained with the fluorescence imaging system or optical sectioning fluorescence microscopy.

### Data quantification

To measure the overall changes, i.e., morphological changes, in organoid units, we monitored their area, perimeter, height, and major and minor axes based on stereomicroscopic images using ImageJ Fiji [27]. In the cases of bead ring, sphere, and spherical shell units, their diameters (*i.e.*, major and minor axes) were obtained by a computation based on an ellipse equivalent to the outline shape. For the bead shape units, the major and minor axes of the outer and inner circumferences were measured, respectively. For the spherical shell units, the major and minor axes of the outer and inner spheres were measured, respectively.

Pearson’s Correlation Coefficient was obtained from the optical sectioning fluorescence microscopy data using ImageJ Fiji’s “Coloc 2” function.

### Data reproducibility

All values are shown as mean ± standard deviation (SD) unless stated otherwise. Each data was obtained from at least three independently reported experiments.

## RESULTS AND DISCUSSION

### Fabrication of organoid units with various geometries

We first selected and fabricated the organoid units with the geometry necessary to reproduce the nephrons and other structures in the human kidney. We confirmed that we had no problems releasing the organoid units with various shapes from the molds, and could fabricate them with constant reproducibility (**Fig. 1a** and **1b**). Organoid units made with MDA-MB-231 cells contracted in size as they matured (**Fig.1c**). Previous studies have also reported this phenomenon, which is partly due to actomyosin contractility [26], typical of highly invasive or motile cell types. Notably, even organoids with complex shapes contracted while maintaining their overall shape (**Fig. 1d**–**1i**). The bead ring-shaped units also showed their contraction without rupture of their respective connecting parts.

**Fig. 1.**
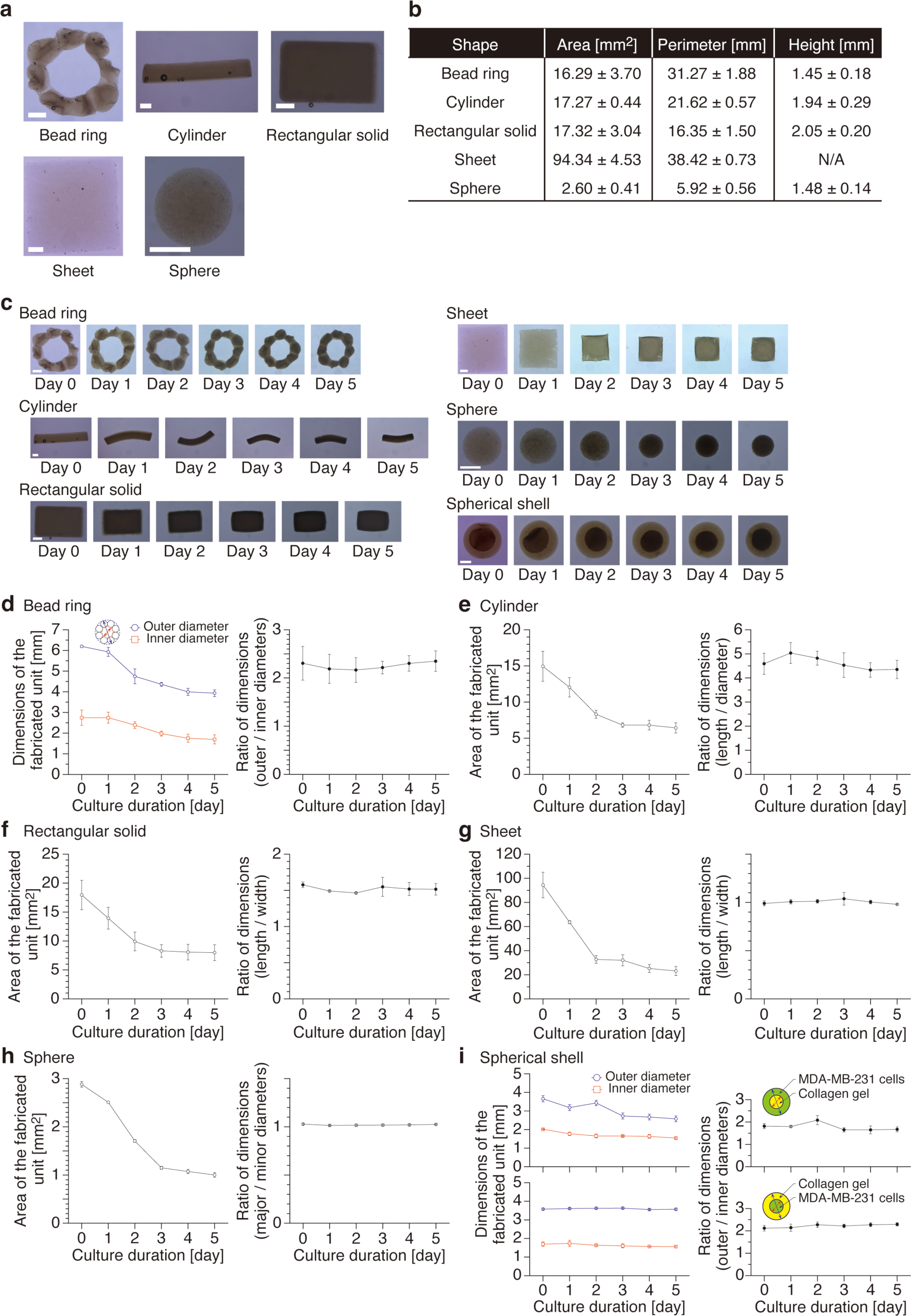
Fabrication of organoid units with various geometries. (**a**) MDA-MB-231 organoid units immediately after fabrication with typical shapes. Scale bars, 1 mm. (**b**) Table of shape reproducibility in organoid fabrication (mean ± SD, *n* = 5). (**c**) Time-sequenced brightfield images of MDA-MB-231 organoid units of various geometries. Scale bars, 1 mm. (**d**–**i**) Dimensional changes of MDA-MB-231 organoid units of various shapes and their ratios with culture duration (**d**, bead ring; **e**, cylinder; **f**, rectangular solid; **g**, sheet; **h**, sphere; and **i**, spherical shell, mean ± SD, *n* = 3). For the spherical shells, we evaluated the case where a layer of cells was placed on the outside and inside, respectively.

Cell proliferation inside and on the surface of the organoids was observed as the culture duration (**Fig. 2a** and **2b**). As the organoid units mature, the inner cells become dense and undergo cell death (**Fig. 2c**). This is caused by the dense cellular area on the organoid surface layer, which acts like a shell and blocks the oxygen and nutrient supply to the inner cells [28, 29]. This is a common problem in previous organoid studies. It can be circumvented by properly arranging supply channels such as the vascular networks [30, 31]. This study allows us to easily fabricate organoid units that reproduce the vascular system as described below. By placing these units inside during assembly, a large-scale organoid that can be cultured for a long period of time is expected to be constructed.

**Fig. 2.**
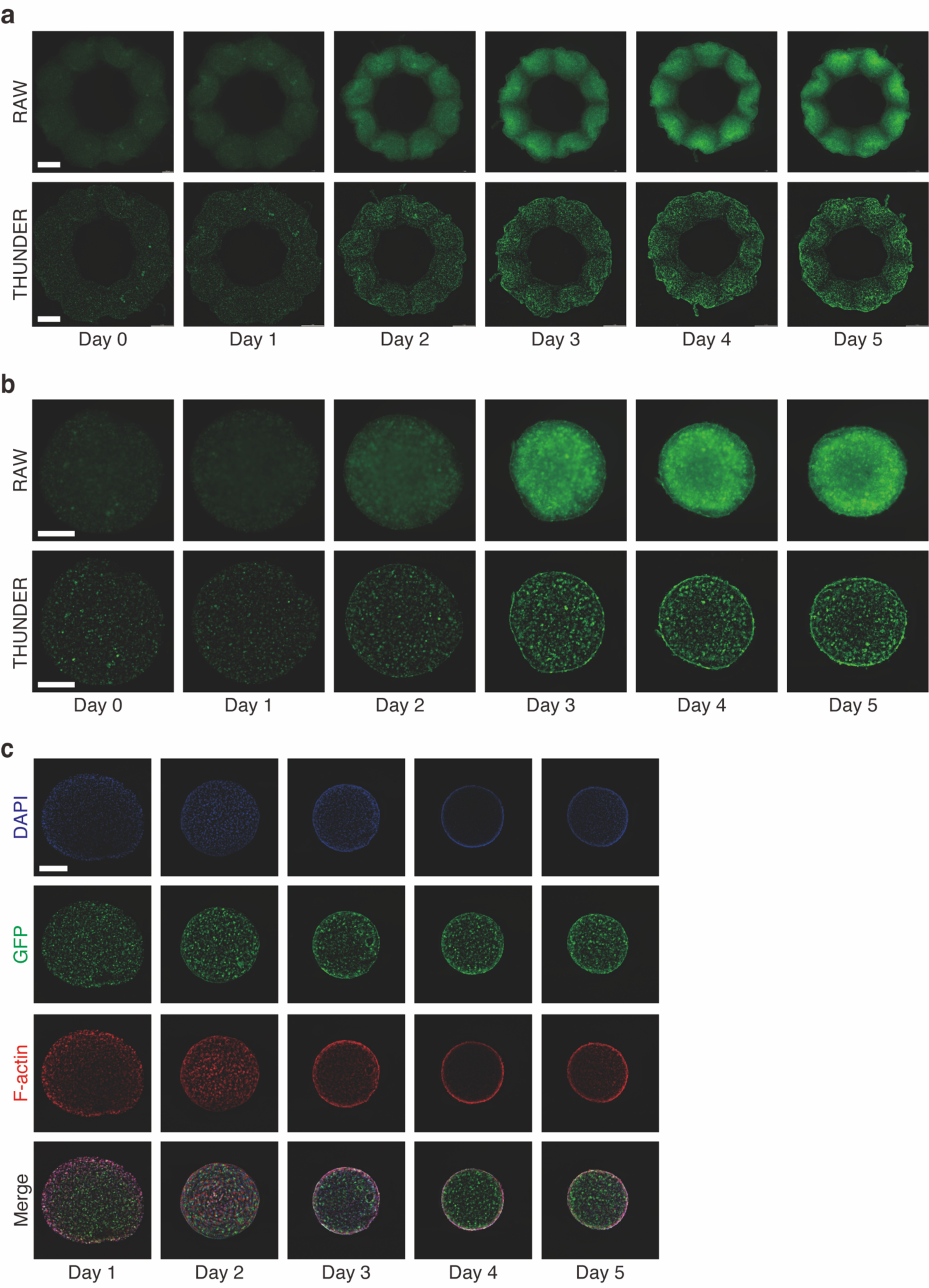
Cell dynamics inside the fabricated organoid units. (**a**) Time-sequenced fluorescence images of a bead ring-shaped MDA-MB-231 organoid unit expressing GFP in the cytoplasm. *Upper row*: wide-field fluorescence images (raw data), *lower row*: images processed by the THUNDER imaging system with instant computational clearing (ICC) and extended depth of field (EDF). Scale bars, 1 mm. (**b**) Time-sequenced fluorescence images of a spherical MDA-MB-231 organoid unit expressing GFP in the cytoplasm. *Upper row*: wide field fluorescence images (raw data), *lower row*: images processed by the THUNDER imaging system with ICC and EDF. Scale bars, 500 µm. (**c**) Fluorescence images of spherical MDA-MB-231 organoid units with stained cell nuclei and the actin cytoskeleton. The images processed by the THUNDER imaging system with ICC and EDF. Scale bars, 1 mm. Quenching of F-actin is observed inside the organoid units around day 4.

We then assembled the various organoid units to evaluate a unit construction method. The substrate, collagen solution, was applied to the area with a micropipette and incubated for 2–5 minutes to allow the units to bond and stack (**Fig. 3a**). Once the four cylindrical units were jointed together to form a structure, we were able to grab one end of the structure with tweezers and pull it up, holding the structure in place without breaking it apart (**Fig. 3b** and **3c**, **Supplementary Movie 1**). Even rectangular units could be easily glued together (**Fig. 3d**), and more units could be stacked on top of the glued units **(Fig. 3e**). The composite of the stacked units did not collapse when shaken (**Supplementary Movie 2** and **3**). After assembly, continued culture is expected to induce proliferation and organization of cells within the units, leading to crosstalk between the units (**Fig. 3f**). However, there are still some problems with the bonding. In this study, the extracellular matrix solution that acts as an adhesive is applied by a micropipette, which results in a large loss due to the inability to apply the solution in a spot manner. This causes a large displacement in the stacking of multiple units. Thus, it suggests that the realization of micro-spot bonding and precise stacking is necessary for the future construction of organ structures.

**Fig. 3.**
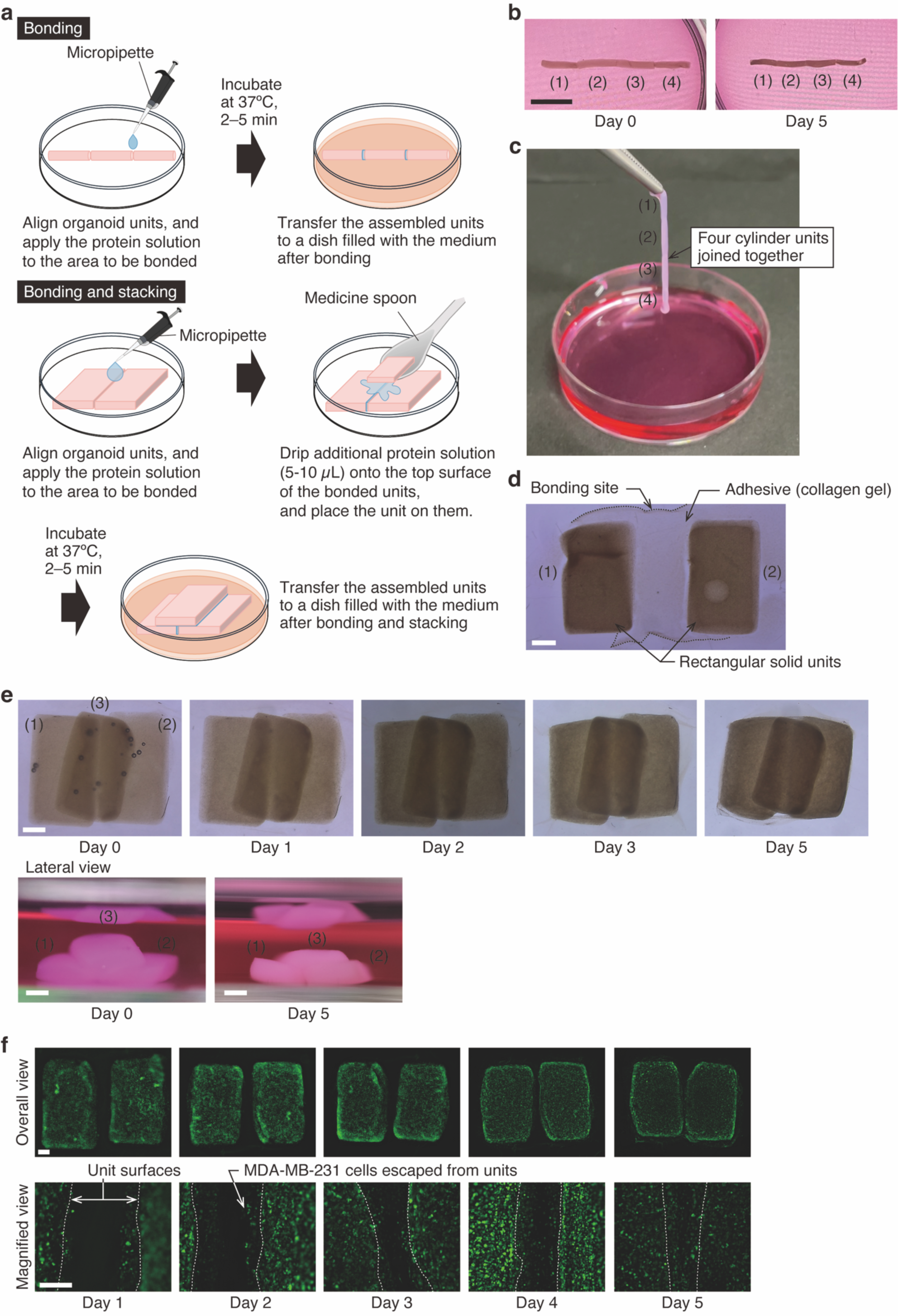
Assembly of organoid units with various geometries. (**a**) Overview of bonding and stacking methods for organoid units. (**b**) Four-cylinder MDA-MB-231 organoid units bonded with collagen solution. Scale bar, 10 mm. (**c**) The jointed units can be pinched and lifted with tweezers without breaking into shreds. (**d**) Two rectangular solid MDA-MB-231 organoid units bonded together. Scale bar, 1 mm. (**e**) Three rectangular solid units were bonded and stacked and showed no signs of dissociation during the five-day culture period. The stacking of the units was confirmed from the lateral view. Scale bar, 1 mm. (**f**) Cellular dynamics around the bonding area between the units (*upper row*: overall view. Scale bar, 1 mm; *lower row*: magnified view. Scale bar, 500 μm). From day 1, cells that escaped from the units began to migrate into the collagen gel at the bonding area. Around day 4, cells proliferated at the bonding area, and the units connect in a cell cluster.

### Construction and assembly of kidney glomerular tissue-like organoid units

The use of easy-to-handle cancer cell lines is suitable for validating the production of living parts of various shapes for full-scale organoid construction. However, it is a considerable leap in research and development steps to construct something that functions as an organ (especially a kidney). Here, we examined the validity of our proposed method (unit construction) by constructing and assembling units with organ-like structures under a tri-culture of parenchymal and interstitial cells (*i.e.*, renal glomerular epithelial cells, mesangial cells, and vascular endothelial cells) that consist of the kidney glomerulus. Tri-cultured organoid units exhibited a complex intertwined tissue-like structure with HUVECs forming a vascular network around HRGEpCs and NHMCs interspersed with them (**Fig. 4a**). These findings are supported by the co-localization of marker proteins of each cell type (*i.e.*, Pearson correlation coefficient; **Fig. 4b**). The internal structure of the produced units is different from the histological structure of human kidney glomeruli [32–34]. The reason for this may be that the tissue did not mature well under the culture conditions in this study due to the lack of orientation of the units. The vascular network formed by the HUVECs decreased with the progression of the culture, and by day 5 the internal network had almost disappeared (**Fig. 4c**). This is most likely due to the fact that the network was recognized as unnecessary because the units were cultured statically. This notion is supported by reports that turnover of the vascular network is induced when the culture medium is not perfused [35, 36]. After the tri-cultured organoid units were bonded to each other with a substrate solution containing interstitial cells (HUVECs), the vascular network formed by the HUVECs seemed to connect the two units (**Fig. 4d**). Crosstalk between units will be possible if the structures with stacked units are cultured under appropriate conditions, including perfusion of the medium, and we can expect the stacked structures to mature as tissues.

**Fig. 4.**
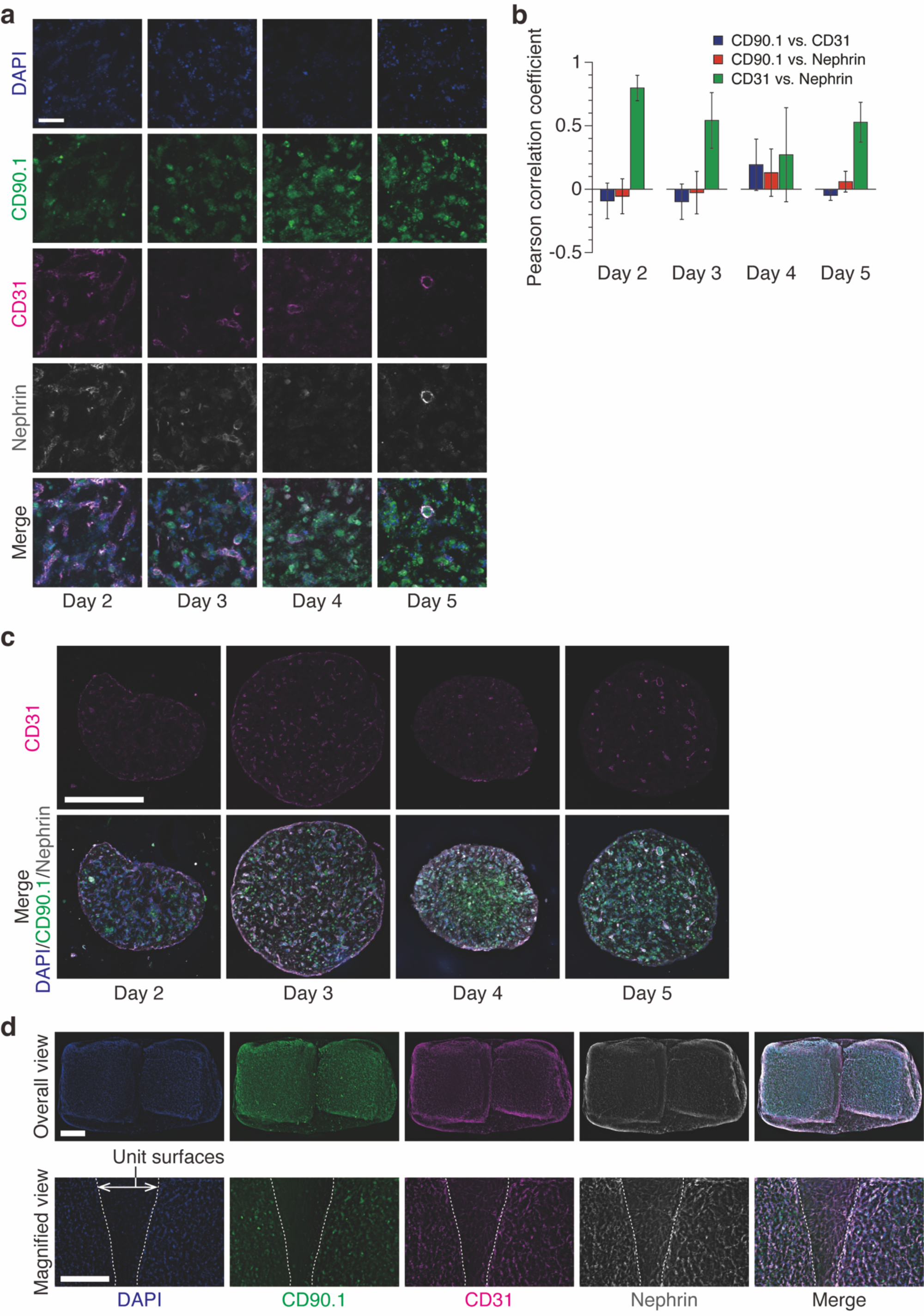
Construction and assembly of human kidney glomerular tissue-like organoid units. (**a**) Fluorescence-stained frozen sections of tri-cultured spherical organoid units. Each marker protein is stained on three cell types: NHMCs (CD90.1), HUVECs (CD31), and HRGEpCs (Nephrin). Scale bar, 50 µm. (**b**) Changes over culture duration in co-localization of marker proteins (Mean ± SD, *n* = 3). (**c**) Overall view of tri-cultured spherical organoid units. Scale bar, 500 µm. (**d**) Cellular dynamics around the bonding area between the tri-cultured cubic organoid units on the second day after bonding (*upper row*: overall view. Scale bar, 1 mm; *lower row*: magnified view. Scale bar, 500 µm).

Our proposed unit construction method can construct tissue-like structures by fabricating and assembling organoid units of various shapes and sizes. Cell viability can also be maintained at a high level by adequately arranging the vascular network that supplies oxygen and nutrients inside under appropriate conditions. Our approach is similar in some respects to the developmental process of tissues and organs because we subdivide the structure of the target organ into elements (units), utilize self-organization of cell clusters at the unit level, and finally assemble them. In contrast, conventional bioprinting technology [21, 37, 38] still needs to solve the issues of bioink development and its influence on cell viability and function, as well as the time required to construct full-scale organoids. In addition, in cases where multiple cell types are used in bioprinting, it is easy to foresee various obstacles to optimizing printing conditions. In general, however, we believe that the greatest challenge for both conventional bioprinting and our method is the expression of higher-order functions in the target organs [39–41] in order to realize the construction of transplantable organoids.

## CONCLUSIONS

In conclusion, we have proposed a method to fabricate and assemble organoid units of various shapes with a size of a few mm as basic elements for the realization of full-scale implantable organoids in the future (**Fig. 5**). Our method is not limited by the ECM types and organoid unit size because it does not use a 3D bioprinter. Moreover, there is no restriction on the adhesive used in our method, so by selecting the appropriate adhesive, it is possible to fabricate large constructions such as assembloids at high speed, which is anticipated to achieve the “highly efficient and high-volume 3D tissue fabrication” required for transplantation medicine. However, it is essential to introduce manipulation techniques such as micromanipulation because precise movements are required to assemble organoid units.

**Fig. 5.**
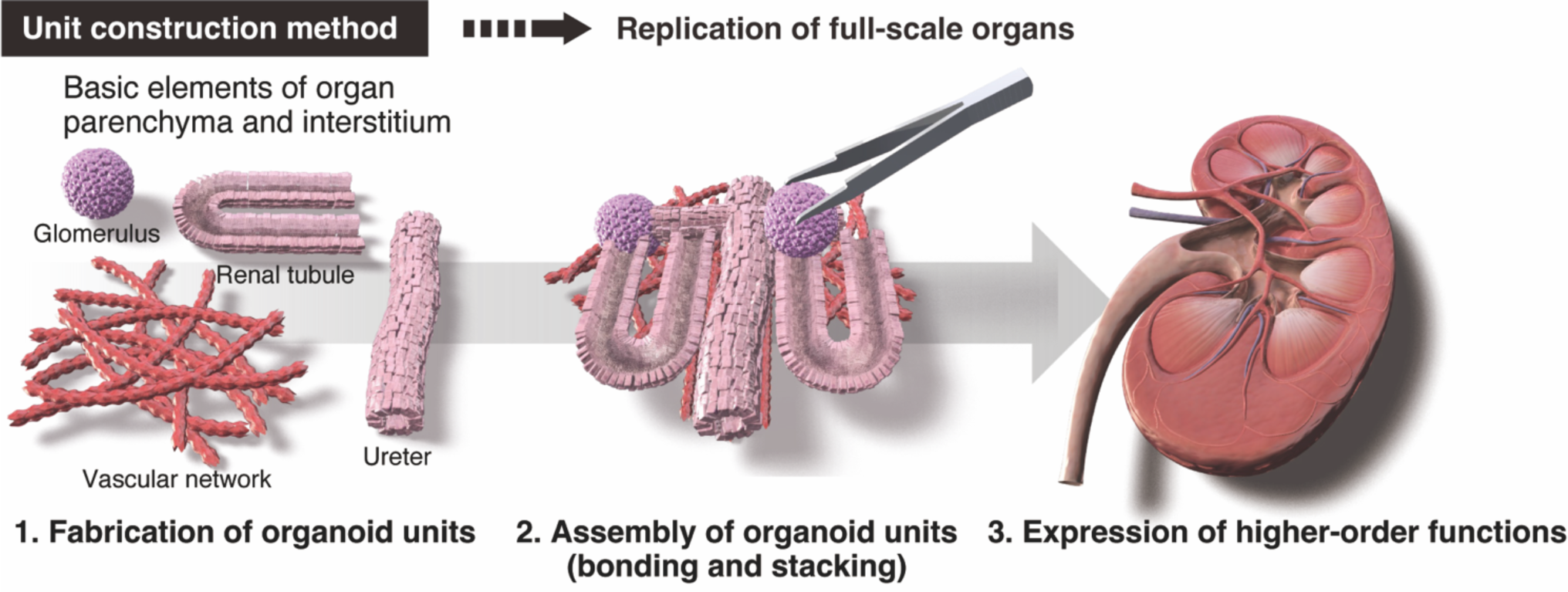
Concept of the proposed unit construction method for replication of full-scale organs with organoids.

## ACKNOWLEDGMENTS

This study was partly supported by grants from the JSPS KAKENHI (Grant No. 21K19893 to D.Y.), the JST FOREST Program (Grant No. JPMJFR222S to D.Y.), and NIMS Joint Research Hub Program (Grant No. 2023-097 to G.H.). The authors would like to thank Science Graphics Co., Ltd. for preparing and editing part of the figures (Fig. 5).

## Conflict of Interest

The authors have no conflicts of interest directly relevant to the content of this article.

## Ethics Approval

Ethics approval is not required.

## Author Contributions

A.K. and D.Y. conceived and designed the research. A.K. conducted most of the experiments. G.H. developed, prepared, and provided the superhydrophobic substrates for organoid block fabrication. All authors discussed the data. A.K. and D.Y. wrote the manuscript. D.Y. directed and supervised the project.

## Supplementary

**Supplementary Fig. 1.**
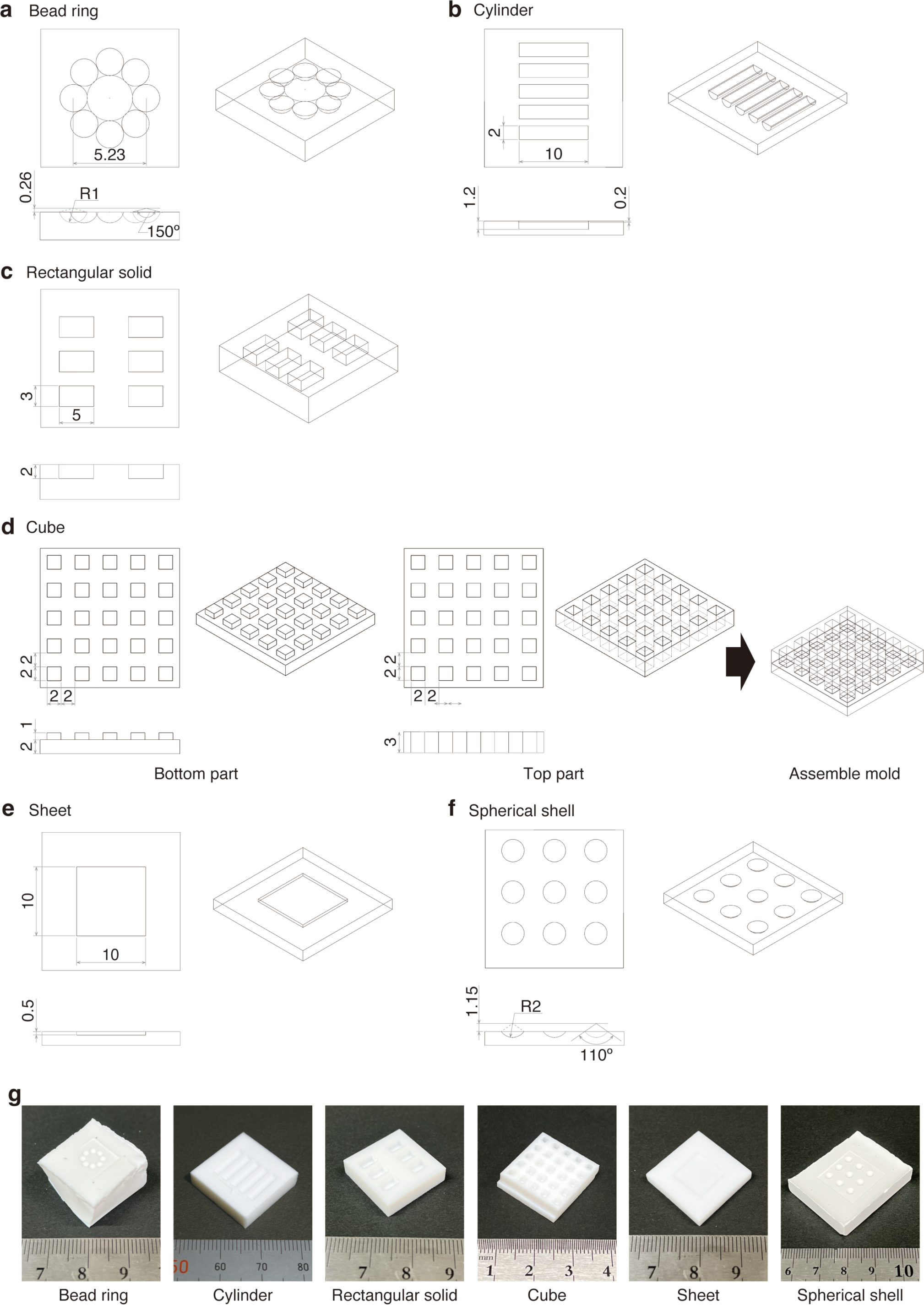
CAD models and actual products of molds to fabricate organoid units with various geometries. (**a**–**f**) Dimensions of various types of molds (**a**, bead ring; **b**, cylinder; **c**, rectangular solid; **d**, cube; **e**, sheet; and **f**, spherical shell). (**g**) Photographs of actual products of the processed molds.

**Supplementary Table 1.**
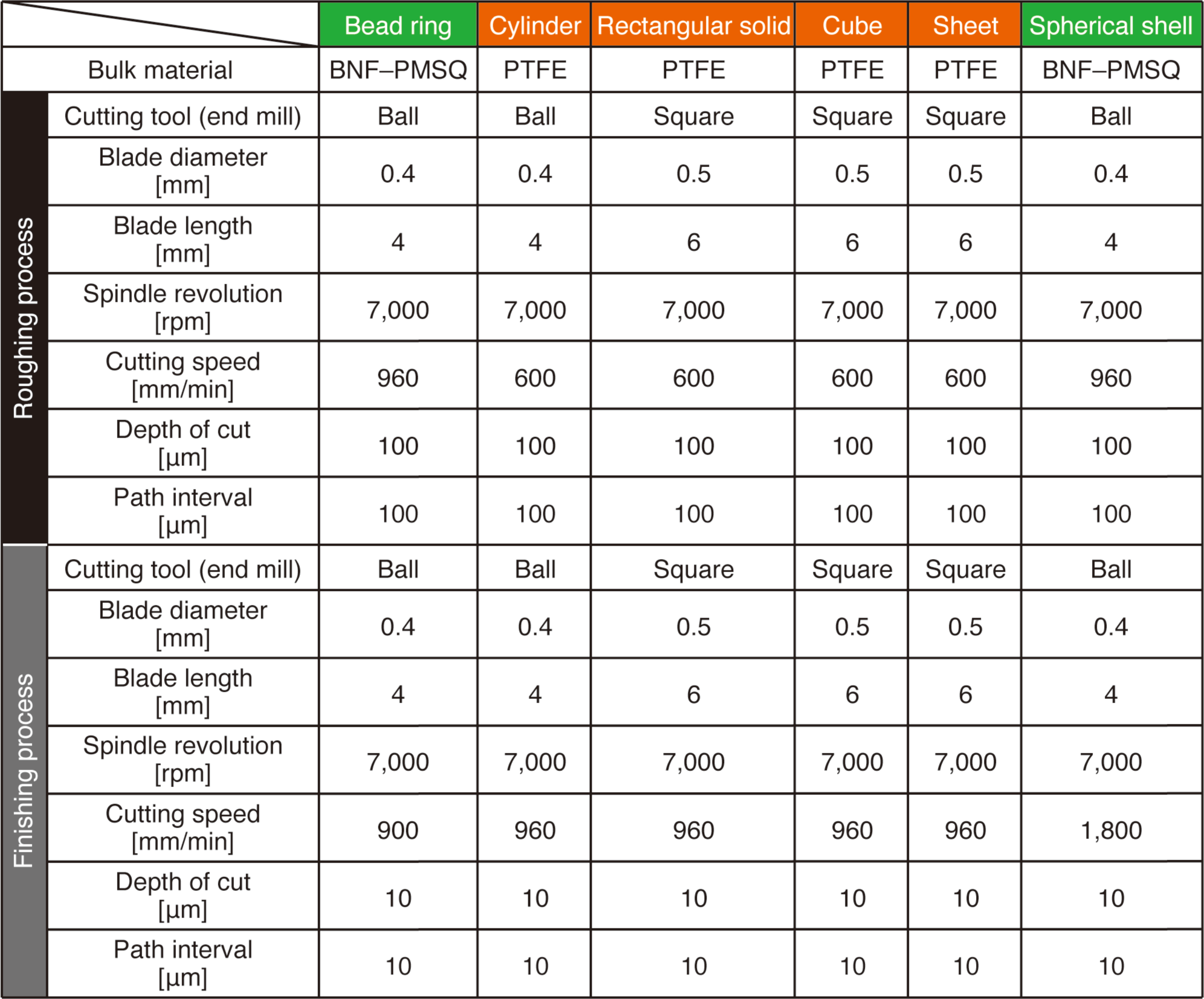
Conditions for milling various types of molds.

## Description of Additional Supplementary Files

**File Name: Supplementary Movie 1**

**Description:** Movie shows lifting a 4-cylinder MDA-MB-231 organoid unit bonded with collagen solution by pinching it up with tweezers.

**File Name: Supplementary Movie 2**

**Description:** Movie shows three rectangular MDA-MB-231 organoid units being shaken immediately after bonding and stacking.

**File Name: Supplementary Movie 3**

**Description:** Movie shows three rectangular MDA-MB-231 organoid units being shaken in five-day culture after bonding and stacking.

## Notes

### Competing Interest Statement

The authors have declared no competing interest.

### Summary of Updates

Figure 3 revised; Figure 5 revised; Supplementary Movie 1 revised.

